# WNT/β-CATENIN modulates the axial identity of ES derived human neural crest

**DOI:** 10.1101/514570

**Authors:** Gustavo A. Gomez, Maneeshi S. Prasad, Man Wong, Rebekah M. Charney, Patrick B. Shelar, Nabjot Sandhu, James O.S. Hackland, Jacqueline C. Hernandez, Alan W. Leung, Martín I. García-Castro

## Abstract

The WNT/β-CATENIN pathway is critical for neural crest (NC) formation. However, the effects of the magnitude of the signal remains poorly defined. Here we evaluate the consequences of WNT magnitude variation in a robust model of human NC formation. This model is based on human embryonic stem cells induced by WNT signaling through the small molecule CHIR9902. In addition to its known effect on NC formation, we find that the WNT signal modulates the anterior-posterior axial identity of NCCs in a dose dependent manner, with low WNT leading to anterior OTX+, HOX-NC, and high WNT leading to posterior OTX−, HOX+ NC. Differentiation tests of posterior NC confirm expected derivatives including posterior specific adrenal derivatives, and display partial capacity to generate anterior ectomesenchymal derivatives. Furthermore, unlike anterior NC, posterior NC transit through a TBXT+/SOX2+ neuromesodermal precursor-like intermediate. Finally, we analyze the contributions of other signaling pathways in posterior NC formation, and suggest a critical role for FGF in survival/proliferation, and a requirement of BMP for NC maturation. As expected RA and FGF are able to modulate HOX expression in the posterior NC, but surprisingly, RA supplementation prohibits anterior, but only reduces, posterior NC formation. This work reveals for the first time that the amplitude of WNT signaling can modulate the axial identity of NC cells in humans.

## INTRODUCTION

Neural crest cells (NCCs) are multipotent embryonic cells that are exclusive to vertebrates, and are endowed with a bewildering differentiation potential enabling them to form peripheral neurons and glia, craniofacial bone and cartilage, and sympathoadrenal secretory cells, amongst other derivatives. During development, expression of early NC markers can be observed above the mesoderm, between the non-neural ectoderm (prospective epidermis) and neural plate (prospective CNS). Then NCCs become transiently incorporated into the dorsal aspect of the neural tube, undergo an epithelial-to-mesenchymal transition, delaminate, and emerge as migratory mesenchymal cells that follow stereotypical paths, and finally differentiate near their final locations. The potential of NCCs to generate terminal derivatives appears to be restricted by their antero-posterior origin, with anterior NCCs contributing to ecto-mesenchymal derivatives, while both anterior and posterior NCCs generate melanoblasts, peripheral neurons, and glia. Given the wide range of derivatives that NC contribute to our bodies, it is not surprising that they are at the root of many taxing pathologies globally known as neurocristopathies.

NCCs are thought to arise through a classic induction event between neural and non-neural ectoderm, and/or mesoderm and ectoderm, reviewed in (Pla and Monsoro-Burq, 2018; Stuhlmiller and García-Castro, 2012a). However, studies in chick and rabbit point to NC specification during gastrulation, and at least for early anterior NC, were shown to be independent of ectodermal and/or mesodermal contribution (Basch et al., 2006; Betters et al., 2018; Patthey et al., 2009). In *Xenopus*, while some studies have also alluded to the induction of NC during gastrulation (Li et al., 2009; Mancilla and Mayor, 1996), a recent study instead proposes that NC are not induced, but instead are a lineage that retains the plasticity and core regulatory mechanisms of pluripotent epiblast/stem cells (Buitrago-Delgado et al., 2015). Overwhelming evidence in various model organisms has supported the participation of multiple signaling pathways during early stages of NC development (Pla and Monsoro-Burq, 2018; Stuhlmiller and García-Castro, 2012a). However, the specific roles of these signals in human NC development remains unresolved, despite their significance to health-related issues.

Efforts to study NC development in human embryos remain limited (Betters et al., 2010; O’Rahilly and Müller, 2007), due to ethical and technical obstacles. However, a powerful alternative emerged through the use of models of NC based on human embryonic stem cells (hESCs) (Pomp et al., 2005). Initial efforts relied on co-cultures, the use of serum or serum replacement cocktails, and either embryoid bodies or neural rosettes (Bajpai et al., 2010; Brokhman et al., 2008; Curchoe et al., 2010; Jiang et al., 2009; Lee et al., 2007; Liu et al., 2012; Rada-Iglesias et al., 2012; Sparks et al., 2018). Simpler models where cell density was controlled along with defined media components have enabled a better analysis of contributing factors during human NC development from hESCs (Fukuta et al., 2014; Hackland et al., 2017; Huang et al., 2016; Leung et al., 2016; Menendez et al., 2011). Common to these and other models of NC development is the activation of the WNT/β-CATENIN signaling pathway (Chambers et al., 2012; Fukuta et al., 2014; Hackland et al., 2017; Leung et al., 2016; Menendez et al., 2011; Mica et al., 2013), which recapitulates WNT’s known role in NC induction in model organisms (García-Castro et al., 2002; LaBonne and Bronner-Fraser, 1998; Lewis et al., 2004).

The WNT/β-CATENIN pathway ultimately results in the accumulation of β-CATENIN in the nucleus where it modulates the expression of targets of the WNT/β-CATENIN pathway through interactions with transcription factors of the TCF/LEF family (reviewed in (de Jaime-Soguero et al., 2018; Nelson and Nusse, 2004). It is noteworthy that WNT/β-CATENIN plays many roles throughout development including early embryonic anterior-posterior axis specification (Loh et al., 2016; Petersen and Reddien, 2009), cellular proliferation (Niehrs and Acebron, 2012), and participates in the formation of neurons, hair cells, mesoderm, and many others cell types (Grigoryan et al., 2008). In hESCs, activation of the WNT/β-CATENIN pathway is known to promote differentiation of endoderm (Naujok et al., 2014), cardiomyocytes (Lian et al., 2012), endothelial cells (Lian et al., 2014), and serotonergic neurons (Kirkeby et al., 2012; Lu et al., 2016), amongst others. Such different effects are thought to be triggered by the specific “context” of the signals and responsive cell, which includes additional transcription factors that cooperate with β-CATENIN, epigenetic signals and modulators, and cross-talk with other signaling pathways (Masuda and Ishitani, 2017).

Other parameters associated with WNT/β-CATENIN signaling such as the magnitude of the signal and duration are also thought to contribute important inputs to the multifactorial process required to deliver specific output responses of the pathway (Goentoro and Kirschner, 2009; Lee et al., 2003; Rogers and Schier, 2011; Tan et al., 2012). Interestingly, while a role for WNT/β-CATENIN signaling in NC development is well established, our understanding of the effects of time and dosage of the signal in neural crest development remain very limited.

In this study, we investigate the effects caused by the magnitude of the WNT signal during human NC formation. We unveil a double bell curve of NC formation mediated by the magnitude of the WNT/β-CATENIN signal, with low and high concentrations leading to anterior and posterior NC formation, respectively. Increasing signal imposes a posterior character in NCCs, as revealed by the induction of posterior identity HOX genes. Interestingly, these posterior NCCs display a mixed differentiation potential, generating peripheral neurons, glia, melanoblasts, in addition to osteoblasts, but unlike anterior NCCs, were not able to differentiate into chondrocytes and adipocytes. Through further interrogation of the formation of posterior NCCs, we find that BMP is required for the generation of posterior NCCs, while FGF signaling appears to regulate proliferation/survival but not the formation of posterior NCCs. Finally, FGF and RA signaling modulate axial character and the expression of HOX genes within posterior NCCs. Together this work demonstrates that modulation of the WNT/β-CATENIN signal promotes the formation of NCCs endowed with different anterior-posterior characteristics.

## RESULTS

### A bimodal wave of NC formation is produced by increasing the magnitude of WNT signaling

We previously reported an effective model of human NC formation that generates anterior NCCs (Leung et al., 2016). In this model, hESCs subjected for 5 days to the WNT agonist CHIR99021 (CHIR) generate >60% cells expressing a wide battery of NC markers, display anterior character, and are able to differentiate into NC-terminal derivatives (Leung et al., 2016). More recently, we explored the effects of temporal variations of the inductive WNT signal on NC formation, and reported that a pulse of CHIR for 2 days, during the start of the differentiation from hESC, leads to a slightly better anterior NC induction rate than the 5-day treatment (G.A.Gomez et al, submitted). To further explore the effect of the magnitude of the WNT signal, we began by assessing the effect of different concentrations of CHIR delivered during the first two days of our NC formation protocol. We initially tested 24 concentrations in 0.5μM increments ranging from 0 to 12μM CHIR, delivered for the first 2 days of the 5 day NC protocol. Fixed cells were then analyzed by immunofluorescence for PAX7 and SOX10. We found high PAX7 and SOX10 expression in cultures receiving 2.5 to 3.5μM CHIR, with reduced levels from 4 to 6μM CHIR, and increasing levels for both markers from 6.5 to 9μM CHIR, followed by sustained high levels of NC markers from 9.5 to 12μM CHIR (Fig. 1A, and data not shown). Tests with 15 μM CHIR displayed no signal for either PAX7 or SOX10 under these conditions, while few cells survived at 20 and 30μM CHIR (data not shown). These results suggest that under our culture conditions, WNT signaling can trigger efficient NC formation from hESCs at a low dosage, near ~3 μM (2.5 to 3.5 μM) and at a high dosage around ~10μM (7 to 12 μM), while intermediate doses between these ranges, or at and above 15μM, do not produce NC.

**Fig. 1.**
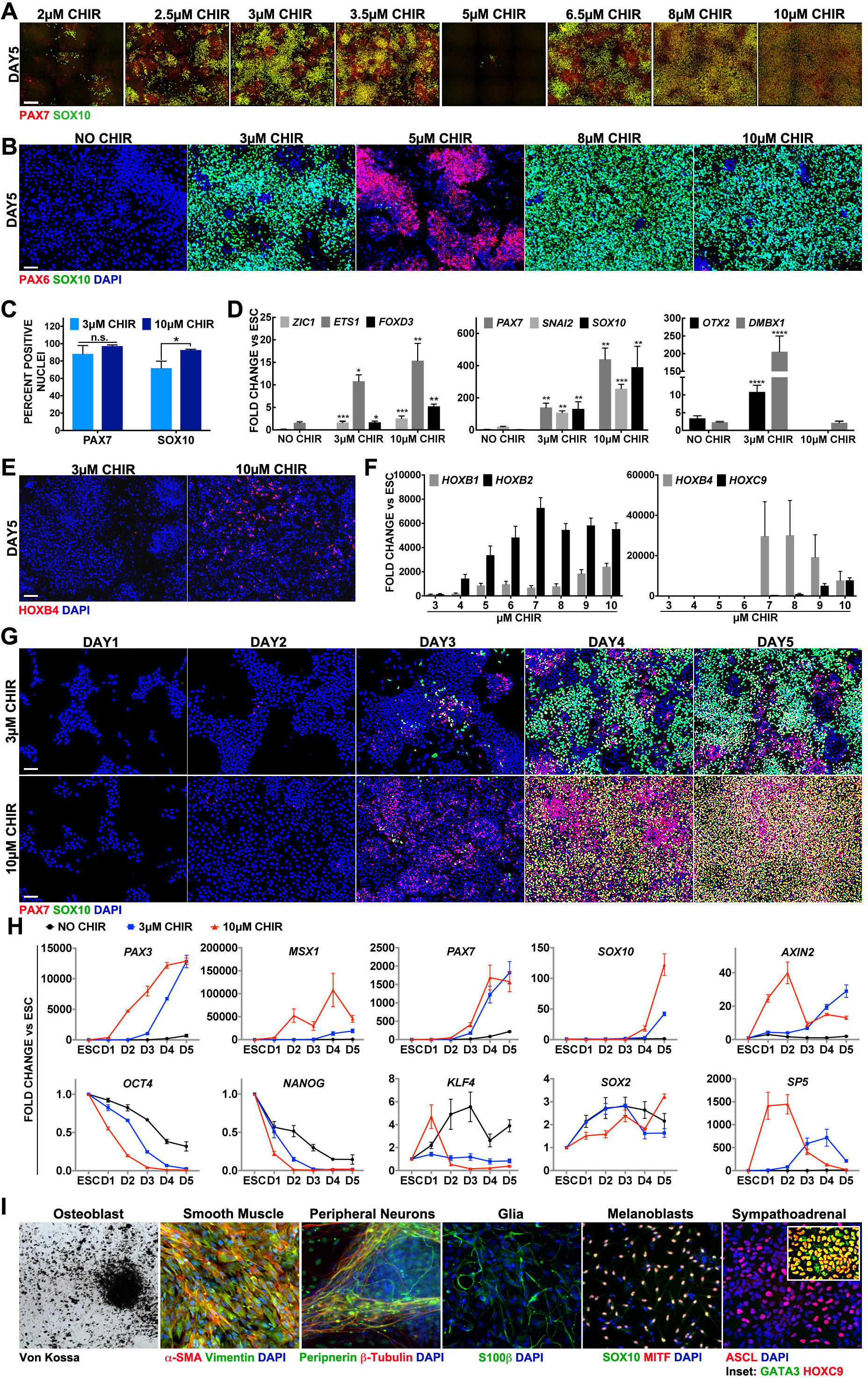
Axial identity of NCCs formed from hESCs is dependent on WNT magnitude, and posterior NCCs retain NC differentiation potential. (A) Immunofluorescence (IF) expression of PAX7 (red) and SOX10 (green) on day 5 after 0-2 day treatment with CHIR at defined concentrations. (B) IF expression of PAX6 (red) and SOX10 (green) on day 5 after 0-2 day treatment with CHIR at defined concentrations. (C) Average percentage of nuclei expressing PAX7 and SOX10 on day 5 by IF. Error bars are ± SEM and *reflects p<0.05 by t-test. (D) RT-qPCR expression levels of NC genes *ZIC1*, *ETS1*, *FOXD3* (left) and *PAX7*, *SNAI2*, *SOX10* (center); and anterior NCs *OTX2*, *DMBX1* (right) on day 5 at CHIR concentrations indicated on the x-axis. Fold change is relative to hESC and normalized by housekeeping genes. Error bars are ± SEM, and data was evaluated by student t-test with each condition tested against the NO CHIR condition * p<0.05 **p<0.05 ***p<0.005 (E) IF expression of HOXB4 (red) at 3μM CHIR and 10μM CHIR. (F) RT-qPCR on day 5 for cultures ranging from 3 thru 10μM CHIR, assessing *HOXB1*, *HOXB2*, *HOXB4* and *HOXC9* with fold change graphed relative to ESCs and normalized by housekeeping genes. Error bars represent ± SEM. (G) IF of time-course images of PAX7 (red) and SOX10 (green) after 0-2 Day treatment with either 3μM CHIR (top row) or 10μM CHIR (bottom row). Each column represents a different day of culture following hESC dissociation and seeding on D0, with a day defined as 24 hours. (H) RT-qPCR expression levels of neural crest markers *PAX3*, *MSX1*, *PAX7, SOX10* (top row, columns 1-4), pluripotency markers *OCT4*, *NANOG*, *KLF4*, *SOX2* (bottom row, columns 1-4) or WNT response markers *AXIN2*, *SP5* (column 5) after NO CHIR (black) 3μM CHIR (blue) or 10μM CHIR (red) treatment on days 0-2 of culture. Fold change is relative to hESC and normalized by housekeeping genes. Error bars are ± SEM. (I) Terminal derivatives obtained from day 5 NCCs induced with 10μM CHIR. Osteoblasts stained with Von Kossa stain,and IF stains for Smooth Muscle, α-SMA (red) & Vimentin (green); Peripheral Neurons, Peripherin (green) & β-Tubulin (red); Glia, S100β (green) & DAPI (blue); Melanoblasts, SOX10 (green) & MITF (red). Nuclei stained with DAPI (blue) where indicated. Scale bars: 225μm in A, and 100μm in B, E, G.

PAX6 has been associated with neuroectoderm fate determination (Zhang et al., 2010). We previously found that during anterior NC formation, cells do not co-express or require PAX6 (Leung et al., 2016). Similarly, SOX10+ NC generated by the 2-day NC protocol with 3, 8 or 10 μM CHIR do not co-express PAX6; however, robust PAX6 expression is seen following transient administration of 4 through 6μM CHIR (Fig. 1B, and data not shown), suggesting a switch from NCCs to neural ectoderm progenitors between the low and intermediate levels of WNT activity.

To compare the cells generated by low and high WNT activation, we evaluated the percentage of nuclei positively stained for PAX7 and SOX10 at optimal CHIR doses. As reported before, low WNT (3μM CHIR) leads to robust expression of both markers. However, treatment with high WNT (10μM CHIR) results in increments of 10% PAX7+ and 20% SOX10+ nuclei relative to low WNT (3μM CHIR) treatment (Fig. 1C). Reverse transcriptase quantitative PCR (RT-qPCR) analysis of NC genes *PAX7*, *SNAI2*, *ZIC1*, *ETS1*, *FOXD3*, and *SOX10* were more strongly expressed following the high CHIR (10μM) dose, further suggesting more efficient NC formation with increased magnitude of WNT activation (Fig. 1D). Given these results, we refer to CHIR treatments as low and high WNT for 3μM and 10μM respectively throughout the rest of the manuscript.

### The magnitude of WNT signal dictates axial identity of NCCs

Increased levels of WNT/β-CATENIN signaling has been shown to impart a posterior character to multiple cell types (Greco et al., 1996; Kiecker and Niehrs, 2001; Kim et al., 2000; Nordström et al., 2002). Therefore, we examined the axial identity of the NCCs induced by low or high WNT. First, we evaluated expression of genes associated with anterior NC territories *OTX2* and *DMBX1* by RT-qPCR on day 5. Both genes are induced by low WNT treatment, however, both are considerably repressed in cells exposed to high WNT (Fig. 1D). In agreement with our results, elevated WNT signaling has been shown to result in the downregulation of *OTX2* in neural ectoderm (Kudoh et al., 2002; McGrew et al., 1997). Given the possible posteriorizing role of WNT, and the lack of anterior markers *OTX2/DMBX1*, we decided to assess the expression of posterior *HOX* genes. We found heterogenous but definite expression of HOXB4 by immunofluorescence in high, but not low, WNT conditions (Fig. 1E).

To further examine the modulation of posterior character of the NCCs generated by different doses of WNT, we monitored the transcriptional profile of various *HOX* genes, *HOXB1*, *HOXB2*, *HOXB4* and *HOXC9* by RT-qPCR. Expression of relatively more anterior HOX genes, *HOXB1* and *HOXB2* were detected at a concentration of 4μM CHIR with amplitude levels increasing as the dose of CHIR is raised (Fig. 1F). By contrast, *HOXB4* first appeared at 7μM CHIR and gradually diminishes as the concentration of CHIR is raised, while *HOXC9* appears at 8μM CHIR and increases thereafter (Fig. 1F). Thus, the degree of NC posteriorization is dependent on the magnitude of the WNT stimulus.

In agreement with our previous findings, NCCs generated under low WNT treatment (3 μM CHIR) display an anterior NC character (Leung et al., 2016). The down-regulation of anterior genes and up-regulation of posterior *HOX* genes in NCCs generated with WNT shows that an increased dosage of CHIR can posteriorize the NC population. Hereafter we refer to anterior NC as those derived from low WNT treatment and posterior NC as those generated from high WNT treatment.

### The transition from hESC to NC is accelerated in posterior NCCs

To better characterize the NC generated with a high WNT dosage, we next explored their differentiation kinetics. To this end, we induced anterior and posterior NCCs, and analyzed the daily acquisition of PAX7 and SOX10. Few SOX10 positive cells appear from day 3 onwards in both conditions (Fig. 1G). However, PAX7 is expressed more robustly on day 3 in posterior NCCs. RT-qPCR analysis of corresponding conditions reveals that in posterior NCCs, other neural plate border specifier genes, *PAX3* and *MSX1* (Pla and Monsoro-Burq, 2018; Stuhlmiller and García-Castro, 2012a), are expressed above basal levels on day 2, a whole 24 hours before their expression is seen in anterior NCCs (Fig. 1H, top row). Thus, increasing the levels of WNT activation to 10μM CHIR leads to a slightly accelerated acquisition of the NC program, but similar to anterior NCCs, the full NC profile arises after 5 days of differentiation.

We then evaluated whether posterior NCCs exit the pluripotent state at a similar time as anterior NCCs do. Transcriptional time-course analysis of core pluripotency markers *OCT4*, *NANOG*, *KLF4* and *SOX2* was assessed by RT-qPCR. *OCT4* and *NANOG* follow the same downward trend in both populations, but they are repressed faster in posterior NCCs (Fig. 1H, bottom row). By contrast *KLF4* and *SOX2* undergo different trajectories. While *KLF4* remains unchanged in anterior NCCs through the 5 days, a surge of *KLF4* is induced by high WNT on day 1 in posterior NCCs that is diminished below levels seen in hESCs from day 2 onwards. Conversely, *SOX2* expression is increased over 2-fold in anterior NCCs during the first 2 days, while in posterior NCCs, *SOX2* only rises about 1.5-fold. Overall, we find that hESCs respond to higher WNT stimulus by altering the pluripotency status sooner than when treated with the lower WNT stimulus.

To confirm our expectation of elevated canonical WNT activity by high WNT treatment, we monitored the expression of direct targets of β-catenin, *SP5* and *AXIN2* (Jho et al., 2002; Park et al., 2013). The levels of both genes were clearly induced to higher levels during the first 2 days in 10μM over 3μM CHIR treatment. However, at day 3, once CHIR was removed from both conditions, a surprising reversal occurred and stronger levels of *AXIN2* and *SP5* are seen in 3μM CHIR. (Fig. 1H, last column). Interestingly, it is clear that this higher level of WNT responsive targets from day 3 onwards is associated with a lower level of expression of NC-markers in anterior NCCs than posterior NCCs. Therefore, the transient and early high dosage of CHIR (0-2 days), correlates with robust NC formation, as seen by PAX7 and SOX10 immunofluorescence and RT-QPCR for multiple NC markers, along with the expression of WNT response targets. Together these results suggest that a transient strong activation of the canonical WNT/β-catenin pathway during the early facet of differentiation is critical to the robust formation of posterior NCCs.

### Posterior NC induced with high WNT can differentiate into posterior terminal derivatives, and display a partial capacity to contribute to anterior ectomesenchymal derivatives

A hallmark of NCCs is their ability to generate a wide range of terminal derivatives. We previously reported that anterior NCCs formed with low WNT generate expected NC-terminal derivatives including melanoblasts, peripheral neurons, glia and ectomesenchymal derivatives (Leung et al., 2016) and G.A.Gomez et al. submitted. We therefore evaluated whether posterior NCCs induced with high WNT (10μM CHIR) can similarly produce terminal derivatives expected of NCCs. We tested their potential to generate peripheral neurons, glia, melanoblasts, and sympathoadrenal derivatives. Simultaneously posterior NCCs were also subjected to the differentiation protocols for 4 anterior ectomesenchymal derivatives including smooth muscle, osteoblasts, adipocytes, and chondrocytes, using specific differentiation protocols for each lineage (Leung et al., 2016). Interestingly, NC generated with high WNT are able to differentiate into peripheral neurons, glia, melanoblasts, and sympathoadrenal derivatives, and failed to generate chondrocytes and adipocytes according to their suspected posterior NC character. However, posterior NCCs also generated smooth muscle, and osteoblasts (Fig. 1I). These results demonstrate that, similar to the anterior NCCs generated with low WNT, posterior NCCs are also multipotent. However, while anterior NCCs are unable to produce sympathoadrenal derivatives, and efficiently generate chondrocytes and adipocytes, posterior NCCs fail to generate the latter and readily produce the former.

### Posterior NCCs require a transient pulse of high WNT stimulus and express molecular markers of NMP-like precursors

Having found that the magnitude of WNT/β-CATENIN signaling affects the axial identity of NCCs produced, we tested the effect of different lengths of duration of the high WNT stimulus via 10μM CHIR application on NC formation. To this end, hESCs were exposed from the beginning of the regimen to high WNT for 1, 2, 3, 4, or 5 days, and all were analyzed after 5 days. Immunofluorescence for PAX7 and SOX10 suggest that posterior NCCs arise robustly when CHIR is added for the first 2 days, but to a much lower extent if added for the first 3 days. While some PAX7 expressing cells appear when stimulated by high WNT for 4 days, minimal SOX10 was found after just one day of treatment, or when treated for 4 or 5 days (Fig. 2A). This is further supported by transcriptional output of *PAX3, PAX7*, *FOXD3,* and *SOX10*, which also exhibit the highest expression levels with 2 days of high WNT treatment (Fig. 2B).

**Fig. 2.**
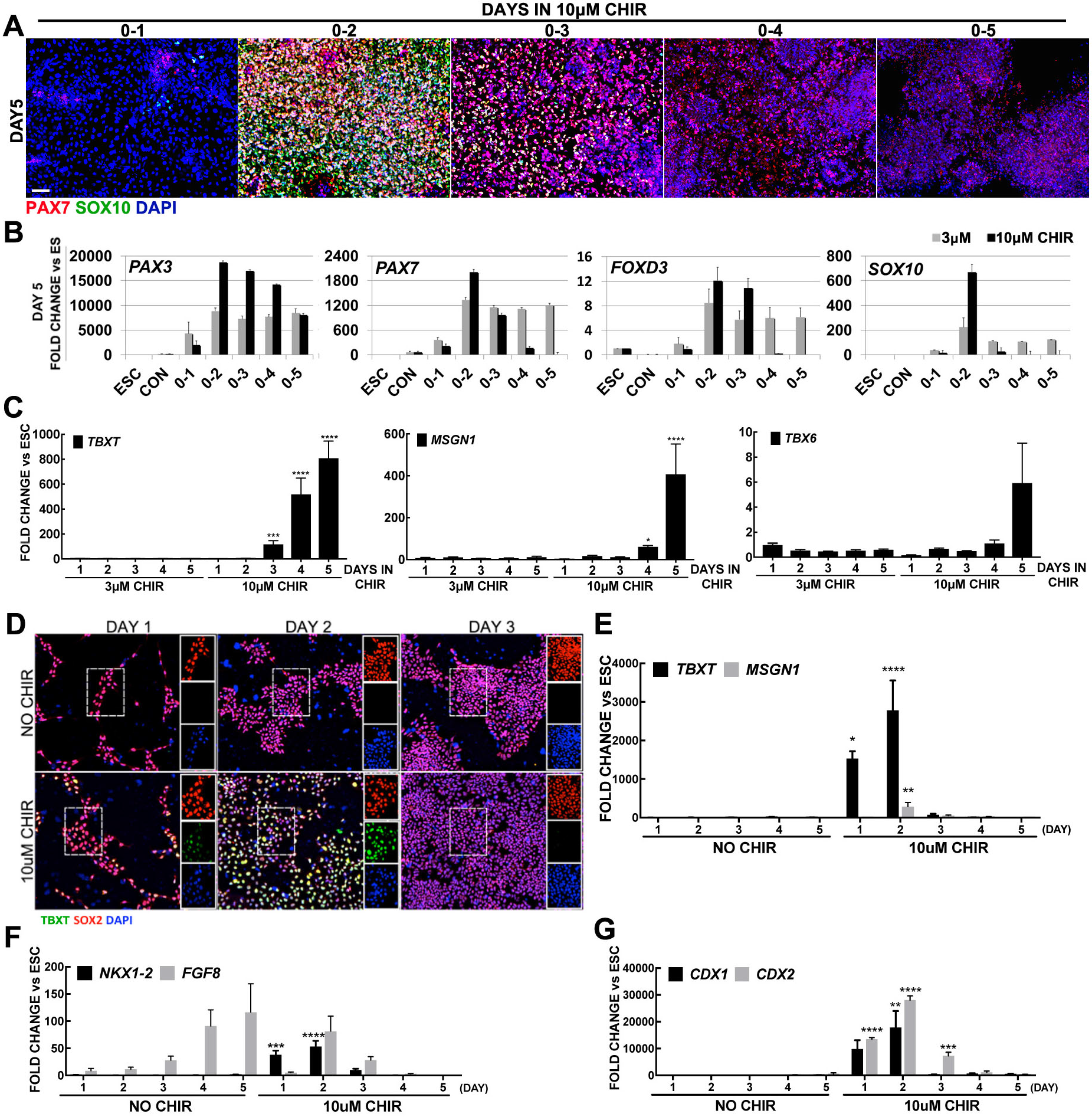
Prolonged treatment with CHIR re-directs the fate of NCCs toward paraxial mesoderm progenitors. (A) IF expression on day 5 of PAX7 (red) and SOX10 (green) following treatment with 10μM CHIR for the duration designated at the top of each image, nuclei were counterstained with DAPI. (B) RT-qPCR evaluation of neural crest markers *PAX3, PAX7*, *FOXD3* and *SOX10* on day5 for NO CHIR controls, CON, and 3μM or 10μM CHIR for durations indicated on the x-axis. Fold change is relative to hESC and normalized by housekeeping genes. (C) RT-qPCR for mesoderm genes *TBXT*, *MSGN1* and *TBX6* on day 5 following treatment with either 3μM or 10μM CHIR for the durations indicated on x-axis. Student t-test was used to compare fold changes between the two doses in matching days of treatment, * p<0.05 ***p<0.005 ****p<0.0005. (D) IF expression of SOX2 (red) and TBXT (green) during the first 3 days in NO CHIR and 2 day treatment with 10μM CHIR. (Sheng et al., 2010) Daily RT-qPCR for genes associated with NMPs, *TBXT*, *MSGN1*, *NKX1-2*, *FGF8, CDX1, CDX2*. Error bars are + SEM, statistical significance was evaluated by t-test between treated and un-treated groups for each corresponding day *p<0.05, **p<0.005, ***p<0.0005, ****p<0.00005. Nuclei stained with DAPI (blue). Scale bar: 100μm

Cultures treated with high WNT for 5 days show minimal PAX7 and no SOX10 protein expression, and no RT-qPCR signal for *PAX7, FOXD3* and *SOX10,* but *PAX3* transcripts were readily detected. Interestingly, PAX3 is known to be strongly expressed and relevant in neural crest and mesoderm development (Epstein et al., 1995; Plouhinec et al., 2014). Therefore, to explore possible mesoderm fate in these cultures, we monitored the expression of other mesodermal markers on a daily basis by RT-qPCR. Cultures treated with low WNT, regardless of the length of exposure, produced no significant signal for *TBXT, MSGN1 and TBX6*, similar to High WNT cultures treated for 1, 2 or 3 days. Instead, treatment with High WNT for 4 or 5 days display robust expression of *TBXT*, *MSGN1* and *TBX6* (Fig. 2C). Therefore, prolonged high-WNT treatment appears prohibitory for the formation of posterior NC and instead triggers the expression of genes associated with a mesodermal fate.

Anterior NCCs produced with low WNT are generated through an intermediate state that does not require nor express PAX6, and while this intermediate progenitor transiently express low levels of *TBXT* transcripts, no protein is detected (Leung et al., 2016). Interestingly, while exploring the ontogeny of high WNT posterior NCCs, we detected strong expression of TBXT mRNA and protein. TBXT is a known target of WNT/β-CATENIN signaling (Yamaguchi et al., 1999), and a prominent mesodermal marker, expressed robustly in the primitive streak, and in posterior neuromesodermal precursors (NMPs), a stem cell population that produces posterior axial tissues including spinal cord and mesoderm (Henrique et al., 2015). A distinct feature of these NMPs is their co-expression of TBXT and SOX2. Evaluation of high WNT cultures at day 2 of treatment revealed NMP features including mRNA expression of *TBXT, MSGN1, NKX1-2, FGF8, CDX1* and *CDX2*, as well as protein co-expression of SOX2 and TBXT (FIG. 2D-G). These results suggest that posterior NCCs generated by high magnitude of WNT transiently express an NMP-like precursor profile.

### BMP signaling is required for induction of posterior NCCs

BMP signaling is also considered a key player in the induction of NC in multiple model organisms (Pla and Monsoro-Burq, 2018; Stuhlmiller and García-Castro, 2012a). Human NC formation models have reported mixed and even contradictory information regarding the contribution of BMP signaling. Some groups require BMP inhibition (Chambers et al., 2012; Mica et al., 2013), but others found BMP inhibition dispensable for NC formation from hPSCs (Fukuta et al., 2014; Huang et al., 2016; Menendez et al., 2011). By contrast, our model of anterior NC formation requires BMP signaling (Leung et al., 2016). We therefore tested whether BMP signaling is also required for posterior NC induction. To this end, we applied the BMP receptor inhibitor DMH1 (Hao et al., 2010) throughout the 5 day posterior NC regimen (2-day high CHIR). Unlike control posterior NC cultures, cells treated with 1μM or 10μM DMH1 do not display PAX7 or SOX10 (Fig. 3A). Interestingly, these cultures reveal a sustained expression of HOXB4 protein by immunofluorescence, suggesting the retention of a posterior character (Fig. 3A). RT-qPCR results also show a reduction in expression of NC markers *PAX7, ETS1, FOXD3,* and *SOX10* in cultures treated with 1 or 10μM DMH1 for 5 days (Fig. 3C).

**Fig. 3.**
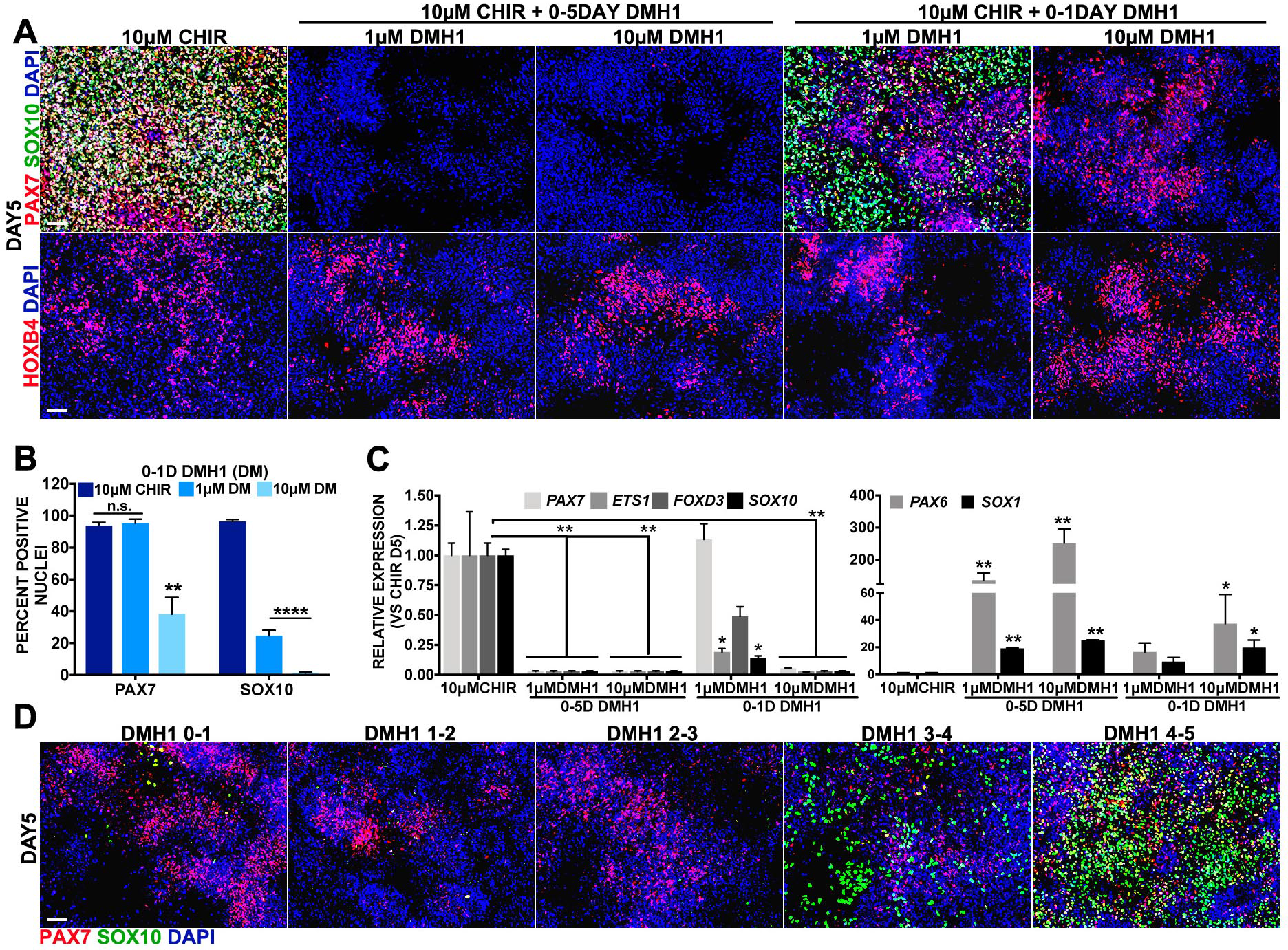
Induction of posterior NCCs requires BMP signaling. (A) IF expressoin of PAX7 (red) and SOX10 (green) in top row, and HOXB4 (red) in bottom row on day 5 after 0-2 day 10μM CHIR treatment; 10μM CHIR plus 0-5 day addition of 1μM DMH1 (column 2) or 10μM DMH1 (column3); or 10μM CHIR plus 0-1 day addition of 1μM DMH1 (column4) or 10μM DMH1 (column5). (B) Average percentage of nuclei expressing PAX7 or SOX10 on day 5 in 10μM CHIR without or with 0-1 day 1μM or 10μM DMH1. Error bars are ± SEM and differences were evaluated by t-test relative to 10μM CHIR, ****p<0.00005. (C) RT-qPCR analysis on day 5 for experimental conditions defined in panel A: 10μM CHIR treatment on days 0-2 and either 1μM or 10μM DMH1 for 0-5 days or on 0-1 day only. NC genes *PAX7*, *ETS1*, *FOXD3*, *SOX10* (left), and neural precursor genes *PAX6* and *SOX1* (right). Data is presented for experimental conditions relative to levels in 10μM CHIR. Errors bars represent ± SEM. Statistics evaluated by one-way ANOVA and changes relative to 10μM CHIR are noted, *p<0.05, **p<0.005 (D) IF expression of PAX7 (red) and SOX10 (green) of cultures analyzed on day 5. All conditions received 10μM CHIR on days 0-2 and addition of 10μM DMH1 on the days indicated above each image. Nuclei stained with DAPI (blue). Scale bars: 100μm

To assess an early requirement of BMP signaling during posterior-NC induction, we deployed DMH1 exclusively for the first 24 hours. DMH1 at a low dose of 1μM for the first 24 hours results in substantially fewer cells expressing SOX10 (Fig. 3A column4, and Fig. 3B), but neither PAX7 or HOXB4 protein were reduced. Similarly, transcripts of *PAX7* are not altered in this condition, even though *ETS1, FOXD3* and *SOX10* are diminished (Fig. 3C). Instead, treatment with 10μM DMH1 during the first 24 hours results in a more prominent inhibition of NC, as determined by reduced immunofluorescence of PAX7 and SOX10 (Fig. 3A column1 vs column5, and Fig. 3B), and by transcriptional repression of *PAX7, ETS1, FOXD3 and SOX10* (Fig. 3C). Concomitant with NC inhibition, in this condition HOXB4 is still expressed (Fig. 3A, last column).

BMP inhibition is a critical instruction during neural development (Chang and Harland, 2007; Linker and Stern, 2004), and is able to redirect anterior NC to neural precursors (Leung et al., 2016). We therefore tested whether posterior-NC cultures were shifted to a neuronal precursor fate upon BMP inhibition by measuring the expression of *PAX6* and *SOX1,* two prominent neuronal precursor genes. Cells treated with DMH1 for 5 days were evaluated by RT-qPCR, and show that compared with minimal expression in 10μM CHIR, both markers are highly induced by BMP inhibition (Fig. 3, right). Conversely, both *PAX6* and *SOX1* are mildly induced by 1 day treatment with 1μM DMH1, but more definitively induced by BMP inhibition with 10μM DMH1. These results suggest that BMP signaling is critical during posterior-NC formation, and its inhibition shifts the fate of the cultures towards a neuronal precursor identity.

We further tested the temporal contribution of BMP signaling to posterior NC formation by interrogating its possible requirement at later time points under the high WNT posterior-NC regimen by delivering the DMH1 inhibitor for 24-hour lapses from days 1 to 2, 2 to 3, 3 to 4, and 4 to 5. Similar to the BMP inhibition on day 0-1, DMH1 delivered on days 1 to 2, and 2 to 3 lead to complete loss of SOX10 expression (Fig. 3D). However, the requirement for BMP was subsequently reduced, since few SOX10 expressing cells emerge when DMH1 was added on days 3 to 4, and a more complete restoration is seen when DMH1 is added on days 4 to 5. Instead, PAX7 expression displays a different dynamic under DMH1 treatment. While cells with BMP inhibition during these later time points display fewer PAX7 cells than controls, their expression is still detectable. These results indicate that BMP signaling is required for posterior NC formation during the first 3 days, and that its inhibition during any 24-hour window during this period is sufficient to prevent the formation of mature posterior NCCs, but not enough to completely prevent the expression of markers associated with earlier stages of NC development, such as PAX7.

### FGF signaling plays a passive role in the induction of posterior NCCs

Together with WNT and BMP, the FGF signaling pathway has also been implicated in NC formation (Betters et al., 2018; Leung et al., 2016; Stuhlmiller and García-Castro, 2012b; Yardley and García-Castro, 2012), and FGF is also known to play an important role in axis specification, particularly contributing to posteriorization (Villanueva et al., 2002). FGF signaling is thought to actively promote early stages of anterior NC specification (Leung et al., 2016; Stuhlmiller and García-Castro, 2012b), but has been proposed to play a passive role in the appearance of posterior NCCs (Martínez-Morales et al., 2011).

To determine whether FGF signaling is required for posterior NC induction, we exposed high WNT cultures to 0.2μM or 2μM of PD173074 (PD17) (Bansal et al., 2003), a potent FGF receptor inhibitor for 5 days. Compared with posterior NCCs generated by CHIR treatment alone, fewer cells survived upon 0.2μM PD17 treatment, but strong PAX7 and SOX10 expression is noted by immunofluorescence in the remaining cells (Fig. 4A). Analysis of immunofluorescence data reveals a mild reduction in the proportion of PAX7 positive (10CH (97.27+1.13%) vs 0-5D PD (91.92+2.16%)) and SOX10 positive (10CH (96.06+1.77%) vs 0-5D PD: (84.72+6.40%)) cells in posterior NCCs inhibited with 0.2μM PD17 (Fig. 4B). Treatment with 2μM PD17 lead to severe reductions in cell numbers that prevented further analysis, although a clear lack of PAX7 and SOX10 expression are noted. RT-qPCR analysis of high WNT cultures exposed to 0.2μM PD17 for 5 days (5D) reveals a 50% reduction in *FOXD3* and *SOX10* and ~30% in *PAX7* and *ETS1* (Fig. 4C, left). To validate the modulation of FGF signaling by PD17, we assessed the expression of known direct targets of FGF, *DUSP6* and *SPRY2* (Ekerot et al., 2008; Minowada et al., 1999). Compared with control posterior NCCs (CH), *DUSP6* and *SPRY2* expression are reduced by 70% and 50% respectively in FGF inhibited cells (Fig. 4C, right).

**Fig. 4.**
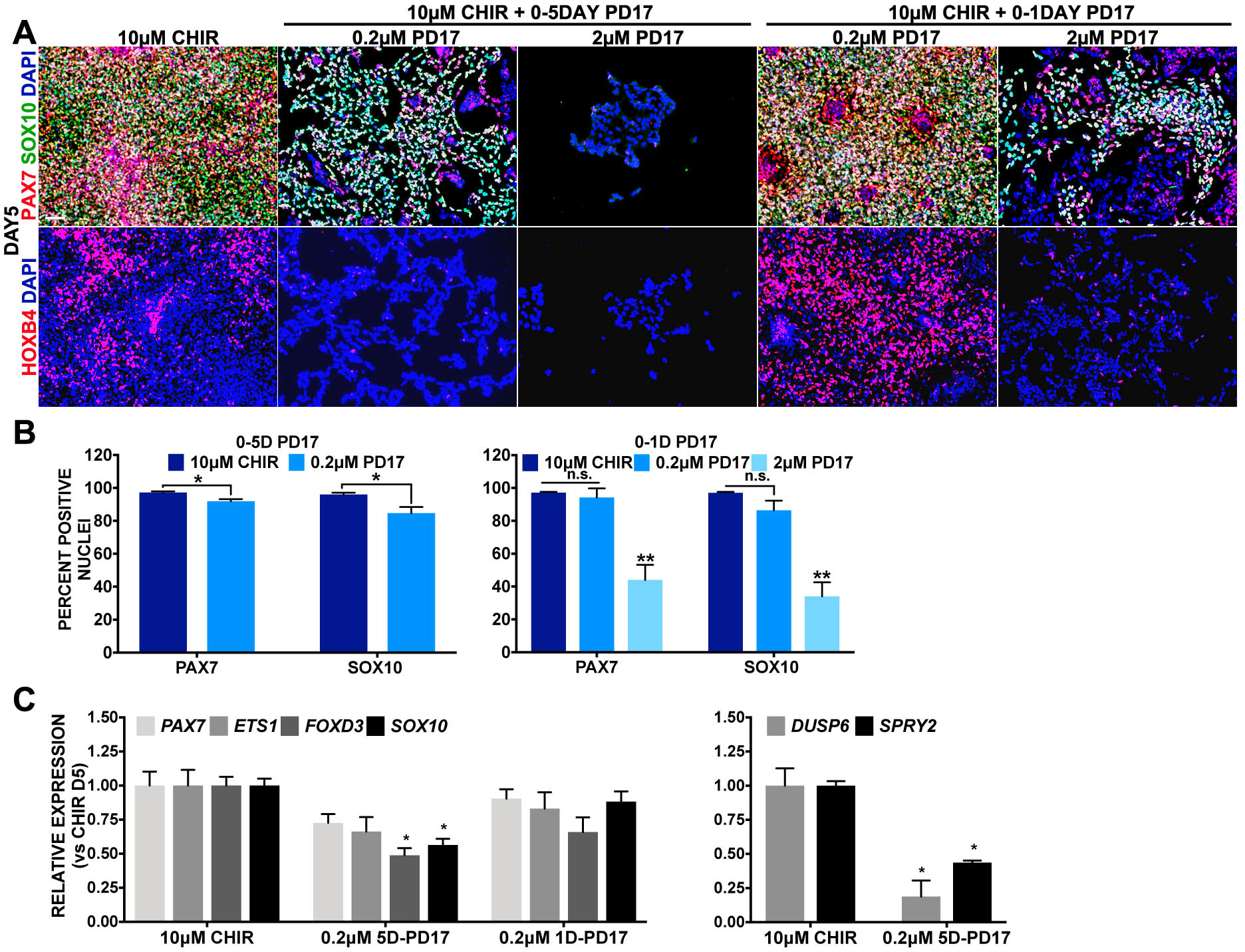
FGF plays a passive role in the induction of posterior NCCs. (A) IF expression of PAX7 (red), SOX10 (green) in top row, and HOXB4 (red) in bottom row, of day 5 cultures treated with 0-2 days 10μM CHIR, or supplemented with 0.2μM PD173074, PD17 (column 2) or 2μM PD17 (column 3) from 0-5 days, and 0.2μM PD17 (column 4) or 2μM PD17 (column 5) on day 1 (0-1D). (B) Average percentage of nuclei expressing PAX7 or SOX10 on day 5 by IF in 10μM CHIR without or with 0-5D PD17 (left) or 0-1D PD17 (right). Error bars are ± SEM and differences were evaluated by t-test, * p<0.05, ** p<0.05. (C) RT-qPCR on day 5 for three conditions, 10μM CHIR, 10μMCHIR plus 0.2μM PD17 from 0-5 days, and 10μM CHIR plus 0.2μM PD17 on day 0-1. NC genes evaluated *SOX10*, *PAX7*, *FOXD3*, *ETS1* (left), and FGF response genes *DUSP6* and *SPRY2* (right). Fold change values are represented relative to 10μM CHIR on day 5. Error bars are ± SEM, statistical significance was evaluated by t-test for each condition relative to 10μM CHIR **p<0.005, ***p<0.0005. Nuclei stained with DAPI (blue). Scale bars: 100μm

Since cell survival is compromised by prolonged FGF inhibition, we tested whether inhibiting FGF during the first 24 hours (0-1day PD17) would affect NC induction, as seen with BMP. We treated hESCs with the 2D High WNT regimen and with 0.2μM or 2μM of PD17 during the first day, and analyzed cultures at the end of day 5. Cell viability was not affected by 24-hour treatment with 0.2μM PD17, and the expression of PAX7 and SOX10 remained unchanged relative to 2D High-CHIR controls as determined by immunostaining (Fig. 4A, 4B). RT-qPCR for *PAX7*, *ETS1, FOXD3* and *SOX10* in these cultures further confirms no deleterious effect on NC formation after 24-hour FGF inhibition (Fig. 4C). In contrast, 2μM PD17 treatment during the first day, was again confounded by low cell survival. However, the few surviving cells reveal expression of PAX7 and SOX10 positive cells, although at notably reduced proportions (Fig. 4B, right). Overall, this data suggests that FGF signaling promotes cellular viability during posterior NC formation, but the cell population that survives do acquire NC character.

Given the known role of FGF signaling in axis specification, we tested whether FGF is required to impart posterior character. To that end, we performed immunofluorescence for HOXB4 in posterior NC cultures treated with PD17. FGF inhibition for 5 days with the “mild” dose of 0.2 μM PD17 led to a reduction in cells expressing HOXB4 while the “strong” dose, 2μM PD17, led to a stark loss of HOXB4 expression upon FGF inhibition for both durations tested (Fig. 4A). Instead FGF inhibition with the lower dose of 0.2μM PD17 for the first day lead to an apparent increase in HOXB4 expression. These results suggest that following a mild short inhibition of FGF signaling, high-CHIR induced NCCs maintain posterior character. Instead, a mild long inhibition (0.2uM for 5 days) as well as a short but strong inhibition (2uM for 1 day) of FGF signaling allow NC formation, but repress the expression of posterior character genes like HOXB4.

### Vitamin A (the precursor of RA) is not required for NC induction or posteriorization, while exogenous RA alters axial identity but inhibits NC formation

Retinoic Acid (RA) signaling has been linked with posteriorization of embryonic tissues and neural crest (Fattahi et al., 2016; Fukuta et al., 2014; Huang et al., 2016; Mica et al., 2013); specifically, addition of RA promotes the expression of vagal identity HOX genes in NC cultures. It has been proposed that RA and FGF exert antagonistic effects to modulate the anterior-posterior character of the neural tube and NCCs (Cunningham et al., 2015; Diez del Corral et al., 2003; Olivera-Martinez and Storey, 2007). We therefore examined whether RA signaling modulates NC formation and their axial character in our human model. One approach used to reduce RA signaling has been to deprive cells of vitamin A, a precursor for RA (Kam et al., 2012). The basal induction media used in our NC protocol contains Vitamin A. Therefore, we evaluated whether anterior and posterior NCCs generated via low and high WNT (3 and 10μM CHIR respectively) are altered by the presence or absence of Vitamin A. Both PAX7 and SOX10 are expressed at normal levels and proportions in anterior NCCs in conditions containing (+) or lacking (−) Vitamin A, and a slight but insignificant reduction was noted in NCCs produced by the high CHIR dose (Fig. 5A). Transcriptional readouts for *ETS1* and *SOX10* further confirm that omission of vitamin A does not affect anterior NCCs but results in a slight reduction in posterior NCCs (Fig. 5B, left). Interestingly, Vitamin A depletion had no effect on the expression of HOX genes in anterior NC, but triggered a slight, although not statistically significant, increase of the expression of *HOXB4* and *HOXC9* expression in posterior NCCs. This is suggestive of a slight contribution of Vitamin A to the known role of RA in the promotion of expression of vagal anterior-posterior axis in the posterior NCCs (Fig. 5B, right).

**Fig. 5.**
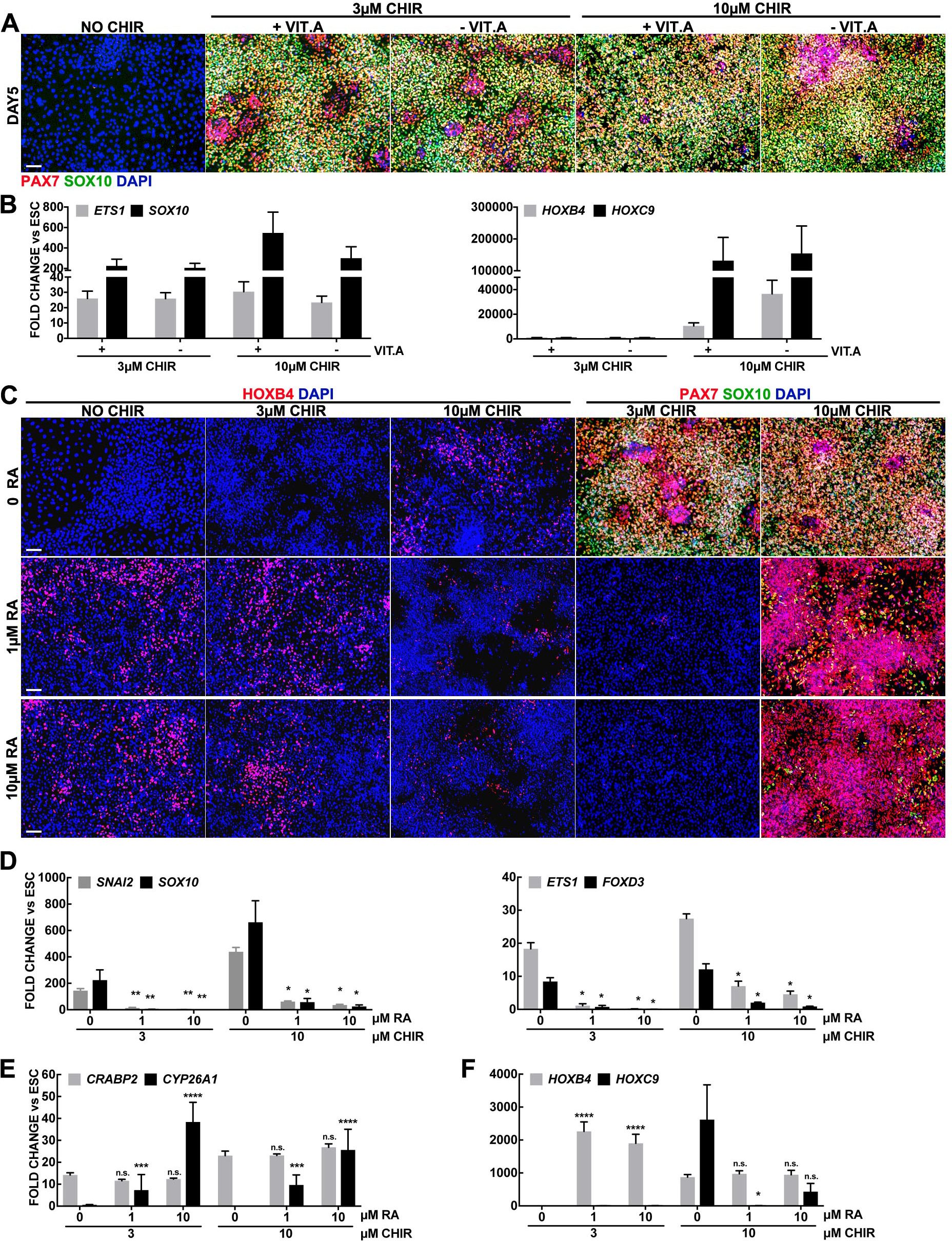
RA is neither necessary for NC induction or posteriorization in high WNT conditions. (A) IF expression of PAX7 (red) and SOX10 (green), on day 5 in basal media containing (+) or lacking (−) Vitamin A (Vit.A). (B) RT-qPCR on day 5 for *SOX10*, *ETS1*, *HOXB4* and *HOXC9* in basal media with (+) or without (−) Vit.A after 0-2 day treatment with 3μM or 10μM CHIR. Fold changes are relative to ESCs and normalized by housekeeping genes. (C)IF expression of HOXB4 (red) in left 3 columns, and PAX7 (red) and SOX10 (green) in right 2 columns on day 5 in cultures treated with NO CHIR, 3μM CHIR or 10μM CHIR on days 0-2 and either DMSO (0 RA), 1μM RA or 10μM RA on days 1-5. (D-F) RT-qPCR on day 5 in 3μM or 10μM CHIR with DMSO (0μM), 1μM, or 10μM RA for (D) NC genes *SNAI2*, *SOX10* (left) and *ETS1*, *FOXD3* (right). (E) RA response genes *CRABP2* and *CYP26A1.* (F) Posterior identity *HOXB4*, *HOXC9*. Fold changes are relative to ESCs and normalized by housekeeping genes. Error bars are ± SEM, statistical significance was evaluated by t-test for each condition relative to either 3μM CHIR or 10μM CHIR *p<0.05, **p<0.005, ***p<0.0005, ****p<0.00005. Nuclei stained with DAPI (blue). Scale bars: 100μm

Next, we tested the role of RA in our model by supplying all-trans-RA (RA) under our culture conditions. To this end, we first assessed conditions upon which RA could induce HOXB4 expression, and found that RA treatment from 1-5 days produced optimal results (data not shown). Next, we added either DMSO (0 RA), 1μM, or 10μM RA on days 1-5 to anterior and posterior NC formation cultures (cells treated for 2 days with 3μM or 10μM CHIR). Surprisingly, we found that the addition of RA to anterior NC cultures blocked NC formation, while instead RA addition to posterior NC cultures appears to increase PAX7 protein while considerably reducing SOX10 expression (Fig. 5C, right). RT-qPCR analysis confirmed that RA treatment down-regulates other NC markers including *SNAI2, ETS1, FOXD3* and *SOX10* in both CHIR treatment conditions, with slightly more resistant expression under the high WNT regimen (Fig. 5D). RA administration induced robust HOXB4 protein expression in NO CHIR, and 3μM CHIR cultures, but in 10μM CHIR cultures a mild reduction of HOXB4 was noted (Fig. 5C, left). By QPCR, *HOXB4* transcripts are elevated in 3μM CHIR condition upon RA treatment, but no evidence of *HOXC9* expression was found. Instead when RA is added to 10μM CHIR cultures, no significant change is seen for *HOXB4,* but *HOXC9* is reduced (Fig. 5F). Confirmation of RA activity is found in the dose dependent up-regulation of the feedback regulator of RA, *CYP26A1* (Loudig et al., 2000) in both 3 and 10μM CHIR conditions (Fig. 5E). However, the expression of *CRABP2*, another putative target of RA (Aström et al., 1992), was not changed under these conditions. Altogether this suggests that activation of RA signaling under these conditions modulates axial character, but antagonizes the NC developmental program.

## DISCUSSION

The WNT/β-CATENIN pathway is known to play a critical role in neural crest formation in multiple model organisms (García-Castro et al., 2002; Saint-Jeannet et al., 1997) and reviewed in (Pla and Monsoro-Burq, 2018; Stuhlmiller and García-Castro, 2012a), and in human models based on PSCs (Chambers et al., 2012; Denham et al., 2015; Fukuta et al., 2014; Hackland et al., 2017; Leung et al., 2016; Menendez et al., 2011; Mica et al., 2013). Simultaneously, this pathway has instrumental roles in fate decisions of many other cell types. The wide range of responses elicited by the WNT/β-CATENIN pathway rely on the availability and sensitivity of a growing number of molecular collaborations that embody the context of the signal. These modulators of specificity include the specific LEF/TCF cofactors, their splice isoforms, other transcription factors, chromatin modifiers, non-coding RNAs and physical parameters such as the initial timing, length of exposure, and concentration of the WNT signal (Goentoro and Kirschner, 2009; Lee et al., 2003; Masuda and Ishitani, 2017; Rogers and Schier, 2011; Tan et al., 2012).

Here we focus on the effect of varying the magnitude of the WNT/β-CATENIN signaling activity on NC formation, aiming to better characterize a key and understudied element of the signaling process. To our knowledge, this is the first time the magnitude of the WNT/β-CATENIN pathway has been addressed in human NC development. We report a bimodal curve of NC formation in response to WNT/β-CATENIN magnitude, with low and high (but not medium or higher) WNT leading to anterior and posterior NC formation. The response of hESCs to WNT magnitude in NC formation and posteriorization provide an intriguing paradigm for modeling the effects of different parameters in the WNT response. It was recently suggested that WNT parameters should modulate the fold levels of β-CATENIN in order to provide distinct effects (Goentoro and Kirschner, 2009). Here we observe a gradual modulation NC posteriorization through gradual HOX gene expression being triggered by relatively small changes in the concentration of the GSK3β inhibitor CHIR, which are difficult to envision as a causative translation to drastic fold changes of β-CATENIN, thus deserving further analysis.

We also report that in our system, posterior NCCs emerge through progenitors that resemble NMPs, noted by co-expression of TBXT+/SOX2+ cells. Furthermore, in this context, BMP signaling is required for both anterior and posterior NC formation, while FGF is required for anterior NC fate acquisition, the posterior NC identity is less dependent on FGF signaling. Finally, FGF and RA contribute with WNT signaling to modulate HOX expression and thus the axial character of posterior NCCs. This study reinforces the importance of the WNT/β-CATENIN signaling pathway in the induction of NCCs in humans and reveal that the parameters of magnitude and duration of exposure to WNT/β-CATENIN not only induces NCCs but can simultaneously alter the axial identity.

### WNT/β-CATENIN promotes NC formation and its magnitude sets their axial identity

The experiments presented here suggest that hESCs can adopt distinct axial NC character guided by WNT parameters. With a low WNT signal generating head/anterior NC formation bearing midbrain markers, and instead, a transient high WNT signal generating gradually more posterior NCCs. These posterior NCCs are derived from TBXT/SOX2 expressing progenitors which acquire progressively more posterior HOX gene expression according to the magnitude of the WNT signal. The robust and immediate induction of CDX factors by high WNT further supports the notion that WNTs set up the axial identity of posterior NCCs, since WNT/β-CATENIN signaling induces CDX expression (Ikeya and Takada, 2001; Sanchez-Ferras et al., 2012; Shimizu et al., 2005) and CDX2 mediates NCC induction via PAX3, MSX1, FOXD3, and potentially other factors (Sanchez-Ferras et al., 2016; Sanchez-Ferras et al., 2012)

The low-WNT response might be related to the known role of WNT/β-CATENIN during early development with initial contribution to axial specification and head formation (Fossat et al., 2011; Heasman et al., 1994; Lemaire et al., 1995; McMahon and Moon, 1989; Smith and Harland, 1991; Sokol et al., 1991). While a second wave of WNT takes part in gastrulation, and displays apparent short gradients from the node/anterior primitive streak (Loh et al., 2016; Morkel et al., 2003; Steinhart and Angers, 2018). We propose that posteriorly, transient WNT levels would be experienced by young NC progenitors, which distance themselves from the WNT source, node/anterior PS, as gastrulation continues in later stages of development.

It is intriguing that SOX10+/PAX7+ NCCs can be formed from pluripotent cells in an apparent bimodal wave with optimal doses of low ~3μM, and high ~10μM CHIR generating anterior (OTX2+/DMBX1+, HOX-) and posterior (OTX2-/DMBX1-, HOX+) NC, with intervening (4-6μM) and higher (>15μM) doses at which NCCs are not produced. At very high CHIR concentrations, few if any cells survived (data not shown). Instead at ~5μM CHIR, healthy cultures display abundant PAX6+ cells, that are negative for PAX7 and SOX10 expression, perhaps indicative of a neural precursor fate. We previously found that expression of PAX6+ cells does not precede the acquisition of NC markers, and that NC is not hampered by PAX6 knock down in our anterior NC model (Leung et al., 2016). Our finding that the low dose of CHIR forms NCCs devoid of HOX expression, while concentrations from 6.5 to 10μM CHIR generates HOX+ posterior NC, unveils a switch between anterior and posterior NC mediated by WNT. Furthermore, WNT signaling is able to gradually regulate the expression of HOX genes in the posterior NC in a concentration dependent manner. The WNT pathway has been shown to play a role in early NC formation (García-Castro et al., 2002; Saint-Jeannet et al., 1997), and is known to modulate the axial identity of neural tissues (McGrew et al., 1995; Michaelidis and Lie, 2008). However, until today, no study had characterized the additional role WNT signaling plays in setting the axial identity of NCCs.

### NC formation and WNT signaling during embryonic development

The bimodal response of hESCs to WNT/β-CATENIN magnitude is intriguing. Here hESCs of presumed identical pluripotency were exposed to different doses of CHIR for the same duration, resulting in distinct populations of NCCs. In the embryo, we assume that anterior and posterior NC arise at different time points, and in distinct locations, enabling the contribution of a wide range of factors to provide a unique context for each NC fate. Still, it seems worthwhile to evaluate if some aspects of a bimodal wave of NC formation triggered by WNT is at play *in vivo*.

Different WNT ligands and receptors are expressed throughout the anterior-posterior body plan, but these are subject to complex regulation that do not translate to homogenous activation of their signals. Expression profiles for some WNT/β-CATENIN targets like AXIN2 and SP5, as well as β-CATENIN reporter models, suggest a weaker early WNT signal anteriorly and a stronger signal posteriorly slightly later, with a gap in WNT activity seen along the anterior-posterior axis for several of these (Harrison et al., 2000; Jho et al., 2002; Maretto et al., 2003; Moro et al., 2012). NCCs are normally considered to arise throughout the entire axis of the neural tube, except from the anterior forebrain. Previous suggestions of gaps in NCCs along the anterior-posterior axis focused on Rhombomeres 3 and 5, pointing to cell death and migratory paths as possible models to explain these gaps. Interestingly, multiple expression patterns for NC markers in diverse studies, offer a distinct gap between the midbrain and the hindbrain signal (Khudyakov and Bronner-Fraser, 2009; Sakai and Wakamatsu, 2005). It is known that NC that have emigrated from the neural tube can move antero-posteriorly at its most dorsal locations (Kulesa and Fraser, 1998), potentially hiding possible areas where gaps in NC formation may exist; further studies are needed to better assess this possibility.

### Putative additional players triggered by CHIR99021

CHIR99021 has been used extensively as a WNT agonist since it has a high affinity for GSK3 kinase, and at the low dose of ~3μM, CHIR is highly specific (Ring et al., 2003). We previously showed that WNT3A or CHIR could both trigger NC development from hPSC, and demonstrated that this process depends on β-CATENIN for NC formation under our low-WNT anterior NC formation conditions (Leung et al., 2016). However, at the high dose of ~10μM a few other kinases may also be inhibited by CHIR (Wagner et al., 2016). Of these, Wagner et. al. reported that 19 are inhibited over 50% of their activity, 6 of which are inhibited more than 70%. However, none of these have been reported to play an active role in NC formation, but whether CHIR promotes NC formation by inhibition of these other kinases will require further investigation. Furthermore, WNT/FZD/LRP6 inhibition of GSK3 is known to trigger at least two other responses independent of the WNT/β-CATENIN pathway, namely the WNT/STOP and WNT/TOR pathways which modify broad protein stability and translation respectively. It is likely that CHIR might trigger similar responses, and thus it is worth evaluating their role in NC formation. In addition, GSK3 kinases have multiple targets and modify various signaling pathways beyond the canonical WNT/β-CATENIN pathway (Acebron et al., 2014; Taelman et al., 2010), and whether any such additional GSK3 targets contribute to human NC formation remains possible and unexplored.

### Effects of other signaling pathways in posterior NC generated via high WNT (CHIR)

In embryos, WNT, FGF and RA signals promote early stages of posteriorization by repressing anterior gene regulatory circuits and simultaneously promoting posterior programs (Andoniadou et al., 2011; Kudoh et al., 2002; Lagutin et al., 2003; McGrew et al., 1997; Shimizu et al., 2006). At the open neural plate regions of embryos undergoing primary neurulation, WNTs and FGFs are the main signaling pathways responsible for neural and mesoderm induction medially (Henrique et al., 2015; Wilson et al., 2009) whereas WNT, and intermediate levels of BMPs, and FGF provide critical inputs for generation of NCs at the neural plate border (Selleck et al., 1998; Stuhlmiller and García-Castro, 2012b; Yardley and García-Castro, 2012), and high levels of BMPs promote ectoderm specification (Wilson and Hemmati-Brivanlou, 1995). In *Xenopus* embryos, intermediate BMP levels are thought to trigger neural plate border formation, which is posteriorized by WNT, FGF and RA signaling pathways, giving rise to prospective NCCs (Villanueva et al., 2002). Human NCCs formed from PSCs have been successfully posteriorized by supplementation with BMPs, FGF and RA (Denham et al.; Fattahi et al., 2016; Fukuta et al., 2014; Huang et al., 2016; Mica et al., 2013). It was therefore important to assess if these other pathways could also modulate the character of NC under high WNT conditions.

The role of BMP signaling in NC development *in vivo* and *in vitro* has proved particularly difficult to tease apart. Gradients providing an intermediate level of BMP signaling have been proposed as critical in *Xenopus* NC formation, however this has not been found in amniotes. Furthermore, some hESC models of NC formation have reported that it is necessary to inhibit BMP signaling during NC formation, while others dispensed with the inhibition of BMP (Chambers et al., 2012; Mica et al., 2013). Instead, we previously reported that BMP signaling is required in our low-WNT anterior NC formation model, and suggested that its function is dispensable for the initial events triggered by WNT, but that it is required thereafter (Leung et al., 2016). Here we report that BMP signaling is also required under the high WNT paradigm of posterior NC formation, since a continuous 5-day BMP inhibition completely blocks NC formation. Instead, mild inhibition with either a low or high concentrations of the BMP inhibitor DMH1 during the first 24 hours leads to moderate and complete blockade of SOX10 expression, but surprisingly has a minimal or mild effect on PAX7 expression. These results support a model in which WNT initiates hESC differentiation towards the NC lineage, and that BMP signaling is required slightly later for the acquisition of more mature traits. This perspective is in alignment with experiments performed with avian embryos (García-Castro et al., 2002; Liem et al., 1997; Patthey et al., 2009; Patthey and Gunhaga, 2014; Selleck et al., 1998; Villanueva et al., 2002).

We previously reported that in the context of low WNT stimulus, FGF inhibition had a deleterious effect on anterior human NC induction (Leung et al., 2016). By contrast, the high WNT regimen which induces posterior NC is only modestly reduced by FGF inhibition, as shown here. Similarly, while FGF inhibition prevents NC formation of anterior NC (Betters et al., 2018; Leung et al., 2016; Stuhlmiller and García-Castro, 2012b; Yardley and García-Castro, 2012), FGF inhibition in posterior regions of chicken embryos does not prevent NC formation (Martínez-Morales et al., 2011), supporting the possibility that there are differences in the role played by FGF in anterior and posterior NC in humans. Our group has developed an independent approach to generate posterior NCCs in which FGF inhibition also fails to prevent NC formation (J.O.S.Hackland et. al. manuscript submitted). This further supports the idea that FGF plays a passive role in posterior NC induction. FGF addition has been shown to upregulate *HOX* genes in NCCs, but this treatment reduced the output of NCCs induced (Mica et al., 2013).

Here we assessed whether RA signaling affects NC formation in our human model. We find that omission of vitamin A has no effect on anterior or posterior NC formation, and *HOX* genes are still expressed in posterior NCCs, indicating that RA is neither required for NC formation, or the posteriorization elicited by WNT signaling. Avian studies have reported that Vitamin A deficiency is required for NC migration, and possibly formation (Martínez-Morales et al., 2011). By contrast, mouse knockouts for one of the RA synthesis enzymes, RALDH2 −/− that display clear defects in posterior identity of cells of the caudal hindbrain, seem to still form NC (Niederreither et al., 2000). NC markers are also expressed in NCCs with pan RAR αβγ KOs (Dupé and Pellerin, 2009), suggesting that RA is not required for the formation of the NC populations studied here. Conversely, while RA supplementation in other systems of NC differentiation from hPSCs have posteriorized NCCs (Fattahi et al., 2016; Fukuta et al., 2014; Huang et al., 2016; Mica et al., 2013), in the system presented here, RA prevents NC formation. A potential reason for these discrepancies includes the possibility that exogenous RA can quench β-CATENIN (Vijayasurian Easwaran and Byers, 1999), but this is context dependent, since in some instances RA and β-CATENIN are co-activators (Liu et al., 2002; Tice et al., 2002). Another possibility is that exogenous RA can modulate FGF (Stavridis et al., 2010), and in conditions where RA activity is elevated, FGF inhibition would prevent activation of the NC program. Since addition of RA to low WNT conditions does not posteriorize NCCs as seen at high doses of WNT alone, it is unlikely that increased magnitude of WNT signaling imparts posterior character to NCCs by activation of endogenous RA.

### Neuromesodermal progenitors and posterior NC

Interestingly, unlike anterior NCCs, which do not express genes associated with mesoderm like TBXT, NCCs induced by high WNT do result in the transient and robust activation of TBXT, in what resembles neuromesodermal progenitor intermediates. Indeed, we find that unlike prospective anterior NCCs, the precursors of posterior NCCs in our culture conditions co-express TBXT and SOX2, as well as *NKX1-2*, *MSGN1*, *FGF8, CDX1*, and *CDX2,* suggesting that they transit through a precursor resembling axial neuro-mesodermal precursors or NMPs (Henrique et al., 2015).

It was recently shown that NMPs produced from hPSCs in vitro have the potential to generate NCCs (Frith et al., 2018), and in a manuscript submitted elsewhere by Hackland et al. we have found that posterior NC generated in a xeno-free (BSA free, and using N2 instead of B27 supplement, and under modulated BMP) platform are also derived from a similar axial progenitor. Some recent evidence from lineage tracing experiments in chick and mice favor the contribution of NMPs to NCCs (Cambray and Wilson, 2007; McGrew et al., 2008; Wymeersch et al., 2016). However, the expression profile of the early NC specifier Pax7 in chick embryos labels the whole posterior open neural plate border, surrounding the NMPs, and thus if these cells were to contribute to the neural folds laterally, they would be joining a neural fold already containing prospective NCCs (Pax7+ cells). Alternatively, the NMPs could contribute to the most posterior region of the neural fold, below the existing Pax7+cells. Future lineage tracing approaches with greater emphasis on studying the contribution of NMPs to NCCs are warranted to address this question.

### Differentiation potential of posterior NC

Our posterior NCCs generate melanoblasts, peripheral neurons and glia, and trunk specific sympathoadrenal lineage derivatives. Simultaneously, our posterior NC also generate ectomesenchymal derivatives which are more often associated with anterior character like osteocytes and smooth muscle (Leung et al., 2016). However, unlike anterior NCCs, posterior NCCs were not able to generate chondrocytes or adipocytes. In multiple *in vitro* experiments, posterior NCCs display unexpected extended potential associated with anterior NCCs (Abzhanov et al., 2003; Dupin et al., 2018; McGonnell and Graham, 2002). It seems that the culture conditions for differentiation reveal a wider potential for NCCs *in vitro* than in *in vivo*, and thus it appears that the inductive signals *in vitro* are more persuasive or stronger than those present *in vivo*. It is therefore critical for future experiments to address minimal signal conditions in culture that effectively differentiate NC into their derivatives according to their axial identity, which might reveal meaningful differences related to *in vivo* potential.

Although forced expression of anterior specific gene networks can drive posterior NCCs to differentiate to anterior restricted ectomesenchymal lineages (Simoes-Costa and Bronner, 2016), in vitro differentiation of ectomesenchymal derivatives from trunk NCCs is possible, but chondrocytes differentiation from trunk NCCs occurs with limited yields (Abzhanov et al., 2003; Dupin et al., 2018).

## CONCLUSION

Here we present evidence implicating the WNT/β-CATENIN pathway as a prime instructor of the NC fate whose magnitude and temporal window of activation can dictate NC axial identity. In this context, BMP operates secondary to WNT signaling, FGF has a role in cell survival and/or proliferation but not NC fate acquisition, and RA is not required for either NC formation or axial identity, while addition of exogenous RA derails NC formation. This study offers insights into the parameters that are relevant for human NC formation, and sheds light on the inputs that produce NCCs with distinct anterior-posterior identity.

## MATERIALS AND METHODS

### RT-qPCR

Total RNA was collected in TRIzol (Thermo Fisher Scientific), cDNA was prepared with Applied Biosystems High Capacity cDNA Reverse Transcription Kit (Fisher Scientific), and RT-qPCR was performed with SYBR Premix Ex Taq II (Clontech) on an Applied Biosystems Step One Plus system. Fold-Changes were evaluated by the ∆∆-Ct method and plotted on Excel or Graphpad prism. qPCR primers used:

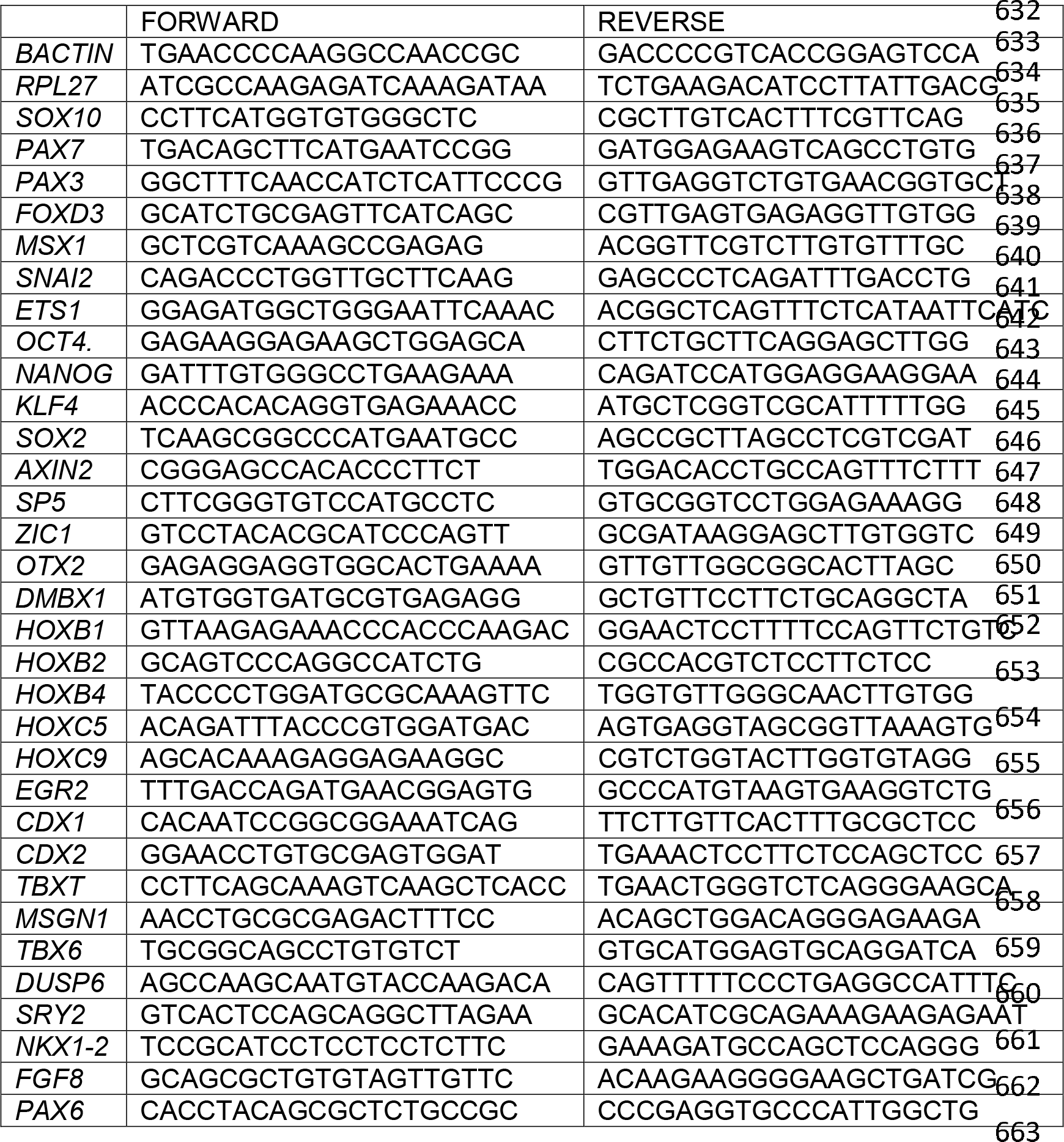

### Immunofluorescence

Cells were fixed in 4% paraformaldehyde, rinsed in PBS, permeabilized with 0.4% triton X100, blocked with 4% fetal bovine serum and incubated with primaries overnight. Alexa Fluor conjugated secondary antibodies (Invitrogen) were used at 1:2000 dilution. Antibodies used: Mouse anti-SOX10 (Santa Cruz Biotechnology, SC271163), 1:200. Mouse anti-PAX7 (1:50), mouse anti-PAX6 (1:100), Rat anti-HOXB4 (1:200) (Developmental Studies Hybridoma Bank). Mouse anti-SMOOTH MUSCLE ACTIN (Sigma, A2547) 1:500. Mouse anti-PERIPHERIN (Santa Cruz, sc-377093), 1:200. Mouse anti- βTUBULIN (Santa Cruz, sc-365791), 1:300. Mouse anti-ASCL1 (Santa Cruz, sc-374104), 1:100. Mouse anti-HOXC9 (Santa Cruz, sc-81100), 1:500. Mouse anti-GATA3 (Santa Cruz, sc-268), 1:100. Mouse anti-VIMENTIN (Sigma-Millpore, MAB3400), 1:1000. Rabbit anti-S100β (abcam, ab52642) 1:200. Goat anti-MITF (R&D systems, AF5769) 1:100.

### Microscopy

Images were captured on a Nikon eclipse 80i microscope on a Spot SE camera and software, or on an inverted Nikon Eclipse Ti microscope with NIS Elements software. All images were compiled and adjusted in Adobe Photoshop CS5.

### Cell Counts

Immunofluorescence images taken on the NIKON eclipse 80i were converted to nd2 format and counted on NIS-Elements AR Analysis software (version 4.6). Nuclei were counted by clustered method at 5px/spot, at a typical diameter of 10μm, with variable contrast levels and dark objects removed.

### Statistics

Statistical analysis of cell counts and RT-qPCR was performed on Prism7.0 (GraphPad Software, San Diego, CA). All experiments were performed with a sample size of n = 3 independent cultures, immunofluorescence images are representative of 1 of these. RT-qPCR was performed from separate wells and 3 technical replicates were assessed for each of 3 independent cultures.

### Neural Crest Differentiation

Human embryonic stem cells line H1 (WA01) obtained from WiCell Research Institute, Inc. (Madison WI, USA), was maintained in mTeSR1 (Stem Cell Technologies) on matrigel coated dishes at 37°C in 5% CO_2_, 5% O_2_ and passaged regularly with Dispase (Stem Cell Technologies) or Versene (Thermo Fisher Scientific). For NC differentiation, hESCs were collected 4 to 5 days after last passage at 80% to 90% confluency. Cells were first rinsed 3 times in 1X PBS Ca+ and Mg+ free (Thermo Scientific, Cat. No. 14190144), then dissociated in Accutase (Stem Cell Technologies) for 4 minutes 30 seconds at 37°C in 5% CO_2_, 5% O_2_. Accutase was immediately quenched with warmed 1X DMEM/F12 (Invitrogen, Cat. No. 17504-044) containing 10μM Rock Inhibitor, Y-27632 (Tocris, Cat. No. 1254), then cells were centrifuged 1200 RPM for 4 minutes at room temperature. 1X DMEM/F12 was discarded leaving cell pellet intact, then replaced with NC Induction media [1X DMEM/F12 (Invitrogen Cat. No. 11320), 1X serum-free B27 supplement (Invitrogen, Cat. No. 17504-044), 1X Glutamax (Thermo fisher scientific, Cat. No. 35050061), 0.5% BSA (Sigma, A7979)] containing 10μM Y-27632. hESC clusters were further dissociated to single cells mechanically by trituration through a 5mL serological pipette a total of 20 times. Cells were counted on a hemocytometer, diluted to an optimal seeding density of 20 × 10^3^ cells/cm^2^ per culture vessel, then seeded on culture vessels pre-coated with Matrigel (BD Matrigel hESC-qualified Matrix Cat. No. 354277). CHIR99021 (Tocris, Cat. No. 4423) was added to cells in induction media during trituration prior to seeding. Note, induction efficiency is sensitive to the CHIR99021 concentration CHIR used, which varies depending on the different lots of CHIR99021, resuspension, pipetting, etc., Optimal inductions in the low WNT were obtained with variable ranges between ~2.5μM - 3μM, but we report all results at 3μM for consistency. 10μM Rock Inhibitor, Y-27632, was added from days 0-2 in all cases and left out of induction media during the remainder of culture. Induction media was changed daily until day of collection for further analysis. Cells were cultured at 37°C in 5% CO_2_, 5% O_2_ throughout the procedure.

### Terminal differentiation of neural crest derivatives

All protocols were adapted and derived from Studer lab group, except chondrocyte differentiation. (Chambers et al., 2012; Lee et al., 2007; Mica et al., 2013) and noted in (G.A.Gomez et al, submitted).

## Acknowledgements

We thank the University of California Riverside, Stem Cell Core for the use of the Nikon eclipse Ti microscope.

## Author Contributions

Conceptualization: G.A.G. and M.I.G.C.; Funding & Resources: M.I.G.C; Performed Experiments: G.A.G.; Experimental support: M.S.P., M.W., J.O.S.H., A.W.L., R.M.C., P.B.S., N.S., J.C.H.; Data Analysis: G.A.G, M.I.G.C.; Writing, review & editing: G.A.G., and M.I.G.C.

## Competing interests

The authors declare no competing financial interests.

## Funding

This research was supported by funding from the National Institutes of Health National Institute of Dental and Craniofacial Research, 5R01DE017914-10

